# Aquatic bacterial metabolic rates from RNA quantitative stable isotope probing (RNA-qSIP) depend on experimental design

**DOI:** 10.64898/2026.07.28.739216

**Authors:** Olivia M. Ahern, Ashley Bulseco, Amy Smith, J.L. Weissman, Joseph J. Vallino, Julie A. Huber

## Abstract

Quantitative stable isotope probing (qSIP) allows researchers to calculate taxon-specific carbon incorporation from sequencing of natural microbial communities, which can be used as a proxy for metabolic activity rates and subsequently as an input for biogeochemical modeling. While qSIP is widely utilized in soils to investigate the identity and metabolic activity of largely unculturable microbes, the application of qSIP in marine and aquatic ecosystems is more recent. Here, we investigated how bioreactor type (batch vs. chemostat) and carbon substrate complexity (single vs. multiple substrates) affect the incorporation of ¹³C-labeled glucose into rRNA after 24 hours using excess atomic fraction (EAF) as a proxy for metabolic activity rate. We found that the growth dynamics and community composition of the ¹³C-incorporating bacteria differed significantly for each treatment. EAF was positively correlated with both 16S gene copy number and a genomic index of copiotrophy in both batch treatments, but not in the chemostat, suggesting that chemostats dampen the competitive advantage of fast-growing copiotrophic taxa. Our results demonstrate that both substrate complexity and experimental regime influence qSIP-derived metabolic activity estimates and provide guidance for future applications of qSIP in aquatic environments.

## Introduction

Earth’s biogeochemical cycles are driven by a complex interplay between individual microorganisms within the framework of geochemical processes. In marine systems, bacteria drive the remineralization of carbon by recycling dissolved organic matter in the ocean (Hansell *et al*., 2009), one of the largest carbon reservoirs on Earth. Despite their importance, quantifying aquatic bacterial contributions to global carbon cycling is challenging due to extremely high taxonomic diversity, low cultivability (Sanz-Sáez *et al*., 2023), and variable growth efficiencies across carbon substrates (Roller and Schmidt, 2015). Uncovering links between bacterial consumers, underlying metabolic mechanisms, and specific consumer-resource dynamics in aquatic systems are key for understanding the carbon cycle and ecosystem health.

Advances in molecular techniques allow for direct measurements of bacterial growth rates *in vitro*. Stable isotope probing (SIP), combined with high throughput sequencing, allows one to track *in situ* microbial activity through incorporation of heavy isotopes into microbial nucleic acids and determine isotope incorporation rates that can be used to infer microbial activity and growth over time (Hungate *et al*., 2015; Purcell *et al*., 2023; Stone *et al*., 2023; Osburn *et al*., 2025). By sequencing the SIP density fractions, researchers can directly identify active bacteria and archaea, their metabolisms, and trophic interactions (Fortunato and Huber, 2016; Wilken *et al*., 2023; Elkassas *et al*., 2025; Coskun *et al*., 2026; Luo *et al*., 2026), without having to isolate microorganisms.

While quantitative SIP (qSIP) can be used to link microbial sequences to species-specific metabolic rates (Hungate *et al*., 2015), experimental artifacts can create bias. A recent soil microbe study demonstrated that both the community composition and assimilation rates derived from DNA-qSIP were significantly different between laboratory and field incubations (Reed *et al*., 2025), making it imperative to understand how similar biases may affect the interpretation of aquatic microbial functions. In aquatic systems, most SIP studies incubate natural communities in batch experiments, where a fixed quantity of labeled substrate is added to the sample in a closed vessel and is tracked. By experimental necessity, the labeled substrate starts at high concentration and can be rapidly consumed over the incubation, which can cause experimental artifacts, especially in experiments focused on heterotrophy. First, carbon amendments in SIP experiments can alter the microbial community being studied. Adding a single labeled carbon substrate introduces a perturbation that may selectively enrich for “copiotrophs” - opportunistic bacteria characterized by rapid growth in high nutrient environments (Zakem *et al*., 2025) - that are not always representative of the *in situ* microbial community (Bulseco *et al*., in press). While the amendment must be high enough to produce a detectable signal in nucleic acids, any carbon addition risks shifting community composition away from *in situ* conditions, making it difficult to determine whether measured metabolic rates reflect natural activity. Second, bacteria *in situ* are exposed to diverse, coexisting carbon sources and experimental evidence shows that bacterial species richness increases in communities grown on media with multiple carbon sources (Enke *et al*., 2019; Dal Bello *et al*., 2021), suggesting single-substrate incubations may not fully capture the full range of community dynamics. Third, in batch growth, both the concentration and growth rate of the added nutrients impact the bacterial growth rate and growth yield (Madigan *et al*., 2008). Consequently, growth rates and incorporation rates measured in single-substrate batch qSIP experiments may poorly represent natural environments, which contain multiple carbon sources and diverse microbial taxa which differ with respect to carbon-use efficiency and carbon specific growth rates (Roller and Schmidt, 2015). Therefore, understanding the limitations of ^13^C-qSIP experimental designs is crucial to determining whether the derived microbial functions reflect those that exist naturally in the environment versus experimental artifacts.

In contrast to batch incubations, chemostats may better mimic natural *in situ* conditions (Gresham and Hong, 2015). In a chemostat, the dilution rate controls the community growth rate and community yield is influenced by nutrient concentrations. Chemostats reach pseudo-steady states with resources maintained at low concentrations that depend on the flow rate into and out of the system relative to its volume (i.e. dilution rate). At steady state, inputs balance losses (Bailey and Ollis, 1986), allowing chemostats to maintain relatively constant cell concentrations and population-level growth. Using complex natural inoculum and conditions mimicking the environment, chemostats can maintain stable biomass while still exhibiting microbial successional patterns (Fernandez-Gonzalez *et al*., 2016; Bulseco *et al*., in press) like those observed in natural ecosystems (Gilbert *et al*., 2012).

Here, we investigated how both 1) batch and chemostat reactors and 2) the availability of either single or multiple carbon substrates, influence microbial metabolic rates and bacterial community composition in SIP experiments. We used excess atomic fraction (EAF) derived from RNA-qSIP as a proxy for *in vitro* metabolic activity rate and calculated two indices for copiotrophy - 16S gene copy number as a proxy for translational efficiency (Roller *et al*., 2016) and an index of copiotrophy (Zakem *et al*., 2025; Weissman *et al*., 2026). We inoculated a complex microbial community (collected from a natural coastal pond community from Cape Cod, Massachusetts) into three different experimental treatments (Figure 1) and followed the incorporation of ^13^C-labeled glucose into complex natural communities after 24 hours. First, we compared EAF and 16S gene copy number in bacteria that incorporated ^13^C-labeled glucose (referred to as incorporators) in single or multi-substrate batch and multi-substrate chemostat experiments. We hypothesized that chemostat conditions would select for incorporators with lower 16S gene copy number and lower index of copiotrophy. Next, we examined differences in EAF from shared incorporators in a batch experiment with a single or multiple carbon sources at equivalent Gibbs Free energy concentrations. We hypothesized that the single-substrate batch incorporators would have higher EAF relative to the multi-substrate incorporators. Determining how experimental design impacts metabolic activity rates is critical to scientists using SIP techniques to understand consumer-resource relationships in aquatic and marine environments.

**Figure 1.**
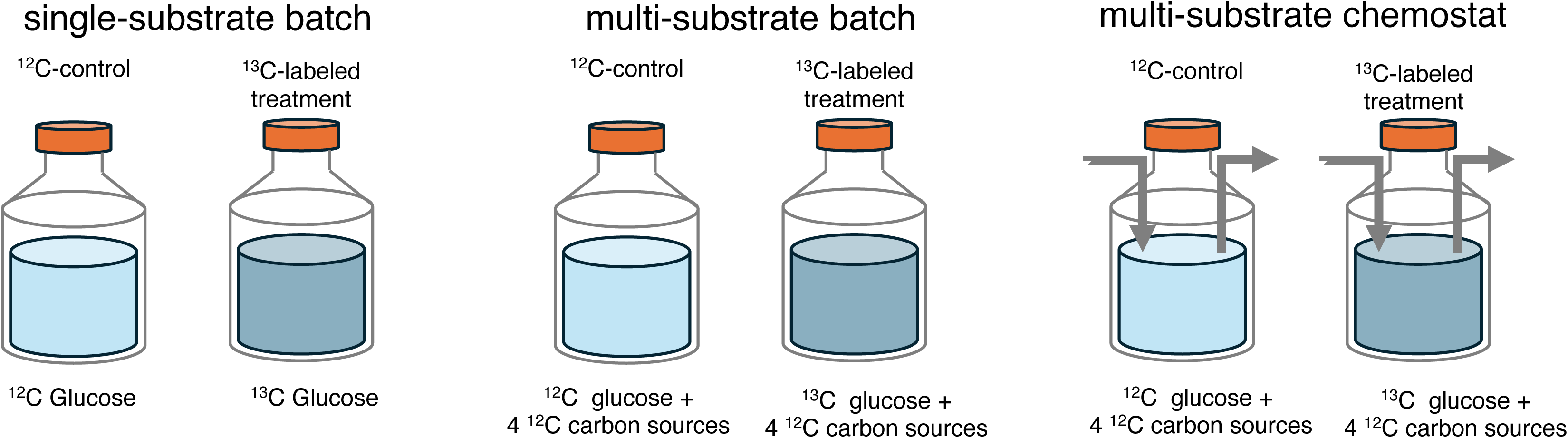
Our experimental design employed aquatic microcosms, amended with either a single carbon substrate (single-substrate) or multiple carbon substrates (multi-substrate), that contained an experimental inoculum with 1356 *μ*M of total carbon at Gibbs free energy equivalent concentrations. ^12^C-controls consisted of either single- or multi-substrates with no isotopically labeled carbon and ^13^C-labeled treatments, where only one carbon source, glucose, was isotopically labeled.

## Methods

### Experimental Overview and Media Design

We ran three carbon amendment treatments using two experimental bioreactor configurations (batch and chemostat) with identical concentrations of salts, nutrients, and trace metals (Figure 1). The three experimental treatments were: 1) <u>single-substrate</u> <u>batch</u>, glucose as the sole carbon source with one ^12^C-control and one ^13^C-labeled treatment; 2) <u>multi-substrate batch</u>, five carbon sources with one carbon source isotopically labeled (glucose) with one ^12^C-control and one ^13^C-labeled treatment; and 3) <u>multi-substrate chemostat</u>, five carbon sources with one carbon source isotopically labeled (glucose) with one ^12^C-control and one ^13^C-labeled treatment (Figure 1). The multi-substrate media contained five carbon sources (206.1 *μ*M methanol, 108.3 *μ*M ethanol, 167.5 *μ*M acetate, 59.69 *μ*M xylose, and 50 *μ*M glucose) with each carbon source added at energetically equivalent concentrations (Bulseco *et al*., in press). The single-substrate media contained 226 *μ*M glucose, energetically equivalent to the total carbon concentration of the multi-substrate media. The ^12^C-control and ^13^C-labeled media had identical compositions, except with the addition of natural abundance or 99.9% ^13^C-labeled glucose (Cambridge Laboratories), respectively.

Experimental media for all of the batch and chemostat contained a defined salt media (30.8 mM NaCl, 2.2 mM Na_2_SO_4_, 0.7 mM KCl, 0.04 mM KBr, 8.0 mM MgCl_2_), 136 *μ*M nitrate, 21.2 *μ*M phosphate, 100 *μ*M CaCl_2_, trace metals (Fernandez-Gonzalez *et al*., 2016; Bulseco *et al*., in press), and were adjusted to ∼pH 7 using a weak NaOH solution and adjusted to 3 PSU with Instant Ocean.

### Experimental inoculum

All experimental treatments were inoculated with surface water collected from a coastal pond (Siders Pond, Falmouth, MA; 41°32’52.08” N, -70°37’26.82” 159 W) on March 6, 2019 for the batch and January 9^th^, 2020 for the chemostat bioreactors. The inoculum water was strained through a 335 *μ*m Nytex Mesh to remove larger particles and organisms.

### Batch

We added 10 % of the Siders Pond experimental inoculum to either single- or multi-substrate media volume of 500 mL in 1L flasks. Flasks were grown in the dark at 25 °C on shaker tables at 150 rpm. Microbial biomass was sampled after 24 hours by filtering 200 mL of the whole community onto 0.22 *μ*m Sterivex (MilliporeSigma) filters, immediately flash frozen in liquid nitrogen and stored -80 °C until RNA extraction.

### Chemostat

Our experimental chemostat setup was similar to those described in (Bulseco *et al*., in press). Three liters of the filtered pond water was added to a 3 L bioreactor (Belco Glass 1964-03600), placed in a dark growth chamber (Conviron PGR15) at 25 °C and sparged with synthetic air at a flow rate of 10 standard cubic centimeters per minute (sccm) from a compressed gas cylinder regulated with a mass flow controller (MKS). The bioreactors were acclimated at these conditions for 5 hours before chemostat operation started.

The chemostat was operated at an initial dilution rate of 0.1 d⁻¹ for ∼24 h to further allow community acclimation, after which the dilution rate was increased to 1.0 d⁻¹ (or 3.0 L d⁻¹ flow rate). The community reached steady state after approximately 19 days, defined by stable dissolved oxygen, pH, and headspace gas concentrations. Once the community reached steady state, the inflowing glucose was switched from ^12^C-control (not isotopically labeled) to ^13^C-labeled glucose, and microbial biomass was sampled after 24 hours. The ^12^C-control corresponded to the microbial biomass sample collected at t = - 0.5, immediately prior to the isotope switch.

Dedicated pumps (Masterflex) were used for the constant inflow of the carbon and nutrient media and the outflow of media from the chemostats to maintain constant volume. To prevent growth in the media, the feeding media was separated into two parts: nutrient only and carbon only. The nutrient media contained 123 *μ*M nitrate and 21.2 *μ*M phosphate, trace metals, and salts to bring it to 3 PSU.

We took both hourly and daily samples from the chemostat. For the hourly measurements, we measured 1) carbon dioxide and oxygen in the headspace via closed gas-sampling loop and 2) dissolved oxygen and pH using controlled and calibrated probes (Bulseco *et al*., in press). Samples were taken daily for 16S rRNA, carbon substrates concentrations, and particulate organic carbon (POC), including its ^13^C percentage (PO^13^C). For RNA sampling, ∼150 mL of sample was filtered onto 0.22 *μ*m Sterivex (Millipore Sigma) filters, immediately flash frozen in liquid nitrogen, and stored - 80 °C until RNA extraction.

We measured the concentration of glucose in the chemostat microcosm through an enzyme assay kit (BIOVISION). Measurements of POC and PO^13^C were calculated at the Marine Biological Laboratory’s Stable Isotope Laboratory.

### RNA extraction, RNA-SIP fractionation, & 16S rRNA library preparation

For all RNA samples, RNA was extracted from the whole Sterivex using the mirVana isolation kit (Ambion) and quantified fluorometrically with RiboGreen (Thermofisher). Gradient preparation, isopycnic centrifugation, and fractionation were performed as described in (Lueders, 2010; Fortunato and Huber, 2016).

To set up our gradient, we combined 500-750 ng of RNA, formamide, cesium trifluoroacetate (Fisher Scientific), and gradient salt solution (Dunford and Neufeld, 2010), to a buoyant density of 1.80 g/mL in a 4.9 mL ultracentrifuge tube (Beckman Coulter). We then spun the tubes for 65 hours at 37,000 rotations per minute at 20 °C in an ultracentrifuge with a VTi 65.2 rotor. Following the completion of the spin, we fractionated our samples into either 12 (batch) or 16 (chemostat) density separated fractions (Dunford and Neufeld, 2010) into 2 mL microcentrifuge tubes (USA Scientific) and purified the RNA with an isopropanol precipitation. We then quantified both the total RNA using RiboGreen (ThermoFisher) and the quantity of the 16S rRNA gene in each fraction using the KAPA SYBR Fast One-Step RT-qPCR kit (Roche) following the manufacturer’s instructions with primers Pro341F and Pro805R (Takahashi *et al*., 2014) for each of the 12 or 16 density separated fractions. Since the density shift between the ^12^C and ^13^C peaks was smaller in the chemostat, we fractionated the chemostat samples into 16 fractions and the batch samples into 12 fractions.

The first strand of DNA was synthesized using the iScript Select cDNA synthesis Kit (Bio-Rad) from the individual RNA fractions using random primers. For both libraries, the 16S V4 rRNA was then amplified in 50 *μ*L reactions using 2 *μ*L of cDNA template, 10 *μ*M 515F (Parada *et al*., 2016) and 806R (Apprill *et al*., 2015) primer set, and GoTaq polymerase and buffer (Promega). The thermocycler program for the PCR was 95 °C for 2 minutes; 30 cycles of 95 °C for 30 seconds, 55 °C for 30 seconds, and 72 °C for 90 seconds; and a final elongation of 72 °C for 5 minutes. The chemostat libraries were sequenced at the University of Connecticut’s Center for Genomic Innovation for MiSeq sequencing with v2 chemistry on a 2x250 bp run and the batch libraries were sequenced at the Bay Paul Center at the Marine Biological Laboratory on a 2x300 bp run (Illumina).

### 16S rRNA bioinformatics, statistics, & identification of bacterial ^13^C-labeled glucose incorporators

All bioinformatic analysis was performed in *qiime2*/2018.11 (Bolyen *et al*., 2019) using *Dada2* (Callahan *et al*., 2016). Unpaired reads were quality controlled, denoised and dereplicated into amplicon sequence variants (ASVs) using *Dada2* with the following specifications: forward and reverse primers were trimmed, filtered to a minimum quality score of 2 and expected error rate of 2, and length of 200 bp and 160 bp for the forward and reverse reads respectively. ASVs were filtered for chimeras using the “consensus” method. ASVs were classified to taxonomy using the V4 region of the Silva 16S database version 132.99 (Quast *et al*., 2013). ASVs were clustered into a midpoint-rooted tree using the *qiime2* script *align-to-tree-mafft-fasttree* using default parameters. We predicted 16S gene copy number using the R package *RasperGade16S* (Gao and Wu, 2023) and normalized libraries by dividing the ASV table by the predicted 16S gene copy number.

All statistical analysis was performed in R version 4.1.2. Sequence tables, trees, taxonomy, and metadata were input into *phyloseq* (McMurdie and Holmes, 2013). ASV tables were filtered for chimeras, mitochondria, chloroplasts, cyanobacteria, and ASVs only assigned to the domain level. Alpha diversity statistics were calculated using *vegan* (Oksanen, 2017). Beta diversity metrics were calculated using philr (Silverman *et al*., 2017).

To identify incorporators (i.e. ASVs that incorporated the ^13^C-labeled glucose), we used quantitative stable isotope probing (qSIP) (Hungate *et al*., 2015) in the HTSSIP R package (Youngblut *et al*., 2018). The qSIP algorithm calculates a quantitative estimate for the percentage of ^13^C atoms in the RNA or excess atomic fraction (EAF), ranging from 0 (natural abundance ^13^C atoms in the RNA) to 1 (RNA is 100% ^13^C atoms). To calculate EAF for each of the ASVs, we normalized the density separated fractions to the concentration of RNA by multiplying the 16S gene copy number normalized reads from each fraction by the number of 16S rRNA copies in each fraction, which was quantified using RT-qPCR. We then filtered the libraries so that one ASV must be present in at least two fractions of both the control and enriched buoyant density gradients. We calculated the EAF for each of the ASVs and excluded EAF values of less than 0.011 since the natural abundance of ^13^C is 1.1%.

We obtained an index of copiotrophy (Zakem *et al*., 2025; Weissman *et al*., 2026) for our incorporators by matching the ASV sequences to the Genome Taxonomy Database (gtdb) version 220 (Parks *et al*., 2022). Representative genomes from GTDB v220 were annotated with prokka v1.14.6 (Seemann, 2014), using option “-kingdom Archaea” for archaeal genomes, and dbcan v 4.1.4 (Zheng *et al*., 2023). We also calculated codon usage bias (dCUB) after filtering organisms with fewer than 10 annotated ribosomal proteins (Weissman *et al*., 2021). Finally, we used the predictIoC function (method="pacific") in the gRodon package to obtain index of copiotrophy values (Weissman *et al*., 2026). Since the index of copiotrophy is strongly phylogenetically conserved up to the genus level (Zakem *et al*., 2025), we averaged the index of copiotrophy for each genera and matched the incorporators which had taxonomy matches to the gtdb genus level.

## Results and Discussion

RNA-qSIP is a powerful tool for linking microbial substrate incorporation and metabolic activity to specific taxa and genes or transcripts. When applied in a quantitative manner, incorporation rates (called EAF or excess atomic fraction) of species-specific metabolic activity rate can be determined without the need for cultivation. RNA-qSIP measures substrate-specific metabolic activity through isotope incorporation into newly synthesized RNA, capturing which taxa are actively metabolizing a given substrate in near real time; this contrasts with DNA-qSIP whose EAF values depend on replication and therefore reflect a more historical, growth-integrated signal. Resolving substrate incorporation rates at the species level moves beyond community-wide measurements to reveal which taxa are responsible for specific biogeochemical transformations. To explore the effects of experimental design on RNA-qSIP metabolic activity rates in aquatic systems, we determined how EAF changed when derived from the same carbon substrate (i.e., glucose; Figure 1) under differing experimental conditions.

Clear separation between the ^12^C-control and the ^13^C-labeled treatment was observed in the all treatments, with a heavier buoyant density peak of rRNA observed in the ^13^C-labeled treatment compared to the ^12^C-control, indicating uptake of ^13^C-labeled glucose into rRNA (Figure S1). The density shift between the rRNA peaks in the ^12^C-controls and ^13^C-labeled treatments was greatest in the single-substrate batch treatment (0.036 g mL⁻¹) compared to the multi-substrate batch (0.017 g mL⁻¹) and multi-substrate chemostat (0.014 g mL⁻¹) treatments (Table 1). The ^12^C peaks for both batch substrate treatments were at 1.77 g/mL while the ^12^C peak for the chemostat was lower, at 1.76 g/mL. While monitoring of the batch experiments was not possible, the chemostat reached a steady state after approximately 15 days (-100 hours before the start of the experiment), indicated by the plateau in chemostat observations for pH, DO, O_2_, and CO_2_ (Figure S2 a,b). Glucose concentrations were near the detection limit in the chemostat experiment while the atomic percent of ^13^C in the particulate organic carbon fraction (> 0.7 *μ*m) reached the expected target of 20 % enrichment after 24 hours (Figure S2c).

**Table 1.** Overview of experimental results with symbol ± indicating standard deviation.

| Description | unit | single-substrate batch | multi-substrate batch | multi-substrate chemostat |
| --- | --- | --- | --- | --- |
| $^{12}\text{C}$ peak | g/mL | 1.77 | 1.77 | 1.76 |
| Differences in $^{12}\text{C}$ and $^{13}\text{C}$ peaks | g/mL | 0.036 | 0.017 | 0.014 |
| No. of $^{13}\text{C}$ glucose incorporators | ASVs | 84 | 60 | 53 |
| EAF | - | $0.34 \pm 0.22$ | $0.27 \pm 0.11$ | $0.15 \pm 0.11$ |
| 16S rRNA gene copy number | copies/cell | $2.75 \pm 1.78$ | $2.63 \pm 1.56$ | $2.55 \pm 1.53$ |
| No. of $^{13}\text{C}$ glucose incorporators matched to index of copiotrophy | ASVs | 62 | 40 | 38 |
| index of copiotrophy | - | $3.22 \pm 0.75$ | $3.23 \pm 0.87$ | $3.69 \pm 0.87$ |

To obtain ASV-specific EAF, we sequenced the 16S rRNA gene of individual density separated fractions from both the ^12^C-control and ^13^C-labeled treatments (Figure S1) and used the qSIP algorithm (Hungate *et al*., 2015) to identify 136 unique ^13^C-labeled glucose incorporators (see Methods). Across all treatments, most incorporators belonged to the bacterial classes Bacteroidia, Alphaproteobacteria, and Gammaproteobacteria. We found that EAF varied among the experimental treatments (Table 1; *anova* F = 21.84, p < 2 × 10 ⁻⁹). We next used excess atomic fraction (EAF), derived from RNA-SIP and as a proxy for ASV-specific metabolic activity rate, to determine if metabolic rates varied due to different experimental conditions. The single-substrate batch incorporators had the highest EAF values (*anova* F = 21.92, p < 2.62 x 10 ^-9^), followed by the multi-substrate batch and chemostat treatments (Table 1). Because growth rate in a chemostat is set by the dilution rate that in turn sets substrate concentration, chemostat EAF likely reflects steady-state metabolic demand rather than opportunistic growth. These results indicate that chemostat SIP experiments may better capture microbial carbon assimilation rates relevant to natural aquatic systems, where substrate supply is typically low and continuous rather than in one large pulse as seen in the batch experiments.

We hypothesized that batch experimental conditions would favor copiotrophic taxa with high growth rates, reflected by high 16S gene copy number, high index of copiotrophy, and high carbon assimilation rates (EAF). The addition of labile carbon can cause the proliferation of high 16S gene copy number bacteria, characteristic of copiotrophs (Klappenbach *et al*., 2000), and DNA-qSIP incorporation rates of ^13^C-labeled carbon were shown to be positively correlated with 16S gene copy number in batch systems (Barnett *et al*., 2021). While mean 16S gene copy number did not differ significantly among treatments (Table 1; *anova*, p > 0.05), we observed positive correlations between EAF and 16S gene copy number in both batch treatments (Figure 2a). EAF and 16S gene copy number were positively correlated for both the single-substrate batch (R^2^ = 0.23, F = 25.92, p = 2.2 x 10^-6^) and multi-substrate batch (R^2^ = 0.049, F = 4.10, p = 0.048) incorporators. We found no correlation between EAF and 16S gene copy number for the multi-substrate chemostat incorporators (p > 0.05). The decoupling of metabolic rate and 16S gene copy number under chemostat conditions suggests that chemostats constrain the advantage of fast-growing, copiotrophic taxa, likely due to the drawdown of substrates to low concentrations. In batch mode, microbes with high 16S gene copy number —and presumably rapid transcriptional capacity—appear to disproportionately assimilate labile carbon, consistent with previous SIP studies (Barnett *et al*., 2021), while the chemostat appears to dampen this “boom–bust” dynamic. Importantly, the lack of correlation between EAF and 16S gene copy number in the chemostat suggests that high 16S gene copy number alone does not confer a competitive advantage under environmentally relevant growth constraints. This finding aligns with observations from natural aquatic systems, where copiotrophic taxa often respond rapidly to carbon pulses but do not dominate under steady-state conditions (Roller and Schmidt, 2015).

**Figure 2.**
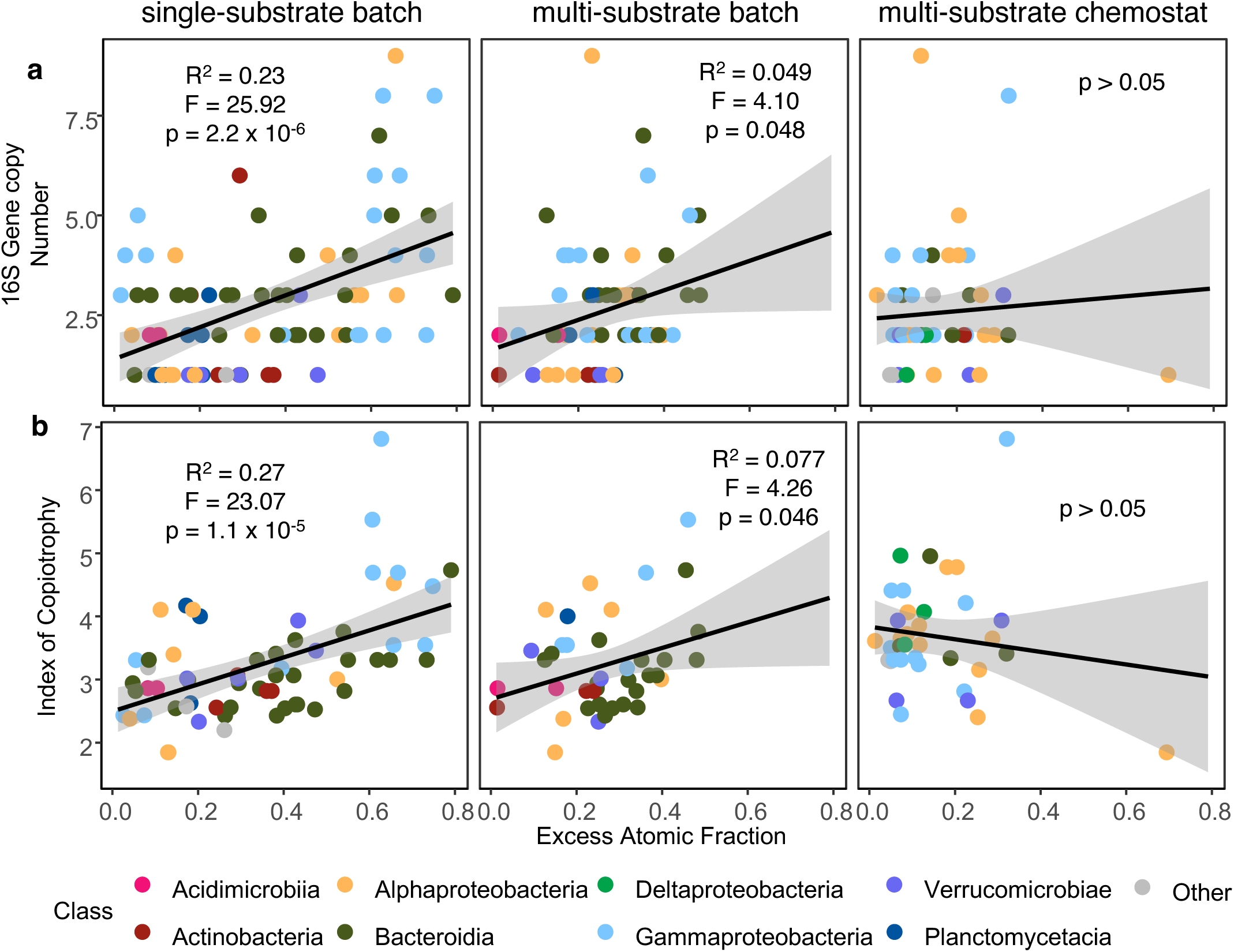
Excess atomic fraction (EAF) versus 16S rRNA gene copy number (a) and index of copiotrophy (b) for the batch and chemostat single- and multi-substrate incorporators. Black lines show linear regression with standard error. Colors indicate bacterial class and “other” indicates bacterial classes with 2 or less incorporators. Correlation coefficients (R^2^), F-value and *p* value are from linear regression.

We next matched our incorporator sequences to the gtdb r220 database to get a genome-based copiotrophic index (Zakem *et al*., 2025; Weissman *et al*., 2026). The gtdb r220 is substantially smaller than the traditional Silva 16S database, therefore only a small portion of our incorporator sequences matched at the gtdb r220 genus level (Table 1). In accordance with the 16S gene copy number results, we found positive correlations between EAF and copiotrophic index in both batch sets of incorporators and no correlation between EAF and copiotrophic index in the multi-substrate chemostat incorporators (Figure 2b). EAF and copiotrophic index were positively correlated for the single-substrate batch (R^2^ = 0.27, F = 23.07, p = 1.1 x 10^-5^) and multi-substrate batch (R^2^ = 0.077, F = 4.26, p = 0.046) incorporators. Therefore, simply being a copiotroph did not always confer a competitive advantage with respect to metabolic activity.

To determine how the incorporation rates of ^13^C-labeled glucose changed due to the complexity of carbon media, we compared the EAF of shared batch incorporators from the single and multi-substrate experiments. We refrained from making direct comparisons at the ASV level between the batch and chemostat for two reasons. First, different CsTFA formulations were used, precluding direct buoyant density comparisons. Second, the batch and chemostat inoculate was collected from different dates from the same sampling location, resulting in statistically distinct community compositions between batch and chemostat experiments (PERMANOVA, R² = 0.50, p < 0.001) making it difficult to distinguish between starting inoculum or selection for differences in community composition between batch and chemostat experimental methods. Therefore, comparisons between the batch and chemostat were made at higher taxonomic levels, EAF values, or predicted 16S gene copy number. In the batch experiments, 57 ASV incorporators were shared between the single- and multi-substrate treatments. Most of these shared incorporators belong to the bacterial classes Bacteroidia (42 ASVs), Alphaproteobacteria (26 ASVs) and Gammaproteobacteria (20 ASVs).

We hypothesized that the EAF of the shared batch incorporators would be higher in the single-substrate than the multi-substrate experimental design because of two potential mechanisms: The first mechanism is that copiotrophic bacteria could outcompete other bacteria in the single-substrate experimental design and therefore incorporate more ^13^C-labeled glucose into their RNA. Another mechanism is the multi-substrate copiotroph’s EAF may be diluted due to consuming several different substrates, or cross feeding on unlabeled or partially-labeled byproducts (Hungate *et al*., 2015; Mooshammer *et al*., 2021). When the EAF values of the shared bacteria in the single substrate experiment were compared to those in the multi-substrate batch experiment, the shared incorporators were positively correlated with EAF value (Figure 3; R^2^ = 0.30, F = 25.38, p = 5.4 x 10^-6^); however, mean EAF was significantly higher for the single-substrate incorporators (*t.test*, t = 3.20, df = 85.18, p-value = 0.002). Diverse assemblages of bacteria can thrive off a single carbon source due to cross feeding (Goldford *et al*., 2018), bacteria are exposed to a variety of carbon sources in the natural environment. Experimental designs with multiple carbon sources increase overall bacterial growth and changing the number of carbon sources decreases competition, allowing for many different species of bacteria to proliferate (Ono *et al*., 2025). While our results suggest that the shared incorporator’s EAF values were positively correlated across the single substrate and multi-substrate bath experiments, the shared incorporator’s EAF derived from ^13^C-labeled glucose did not fall on the one-to-one line (Figure 3).

**Figure 3.**
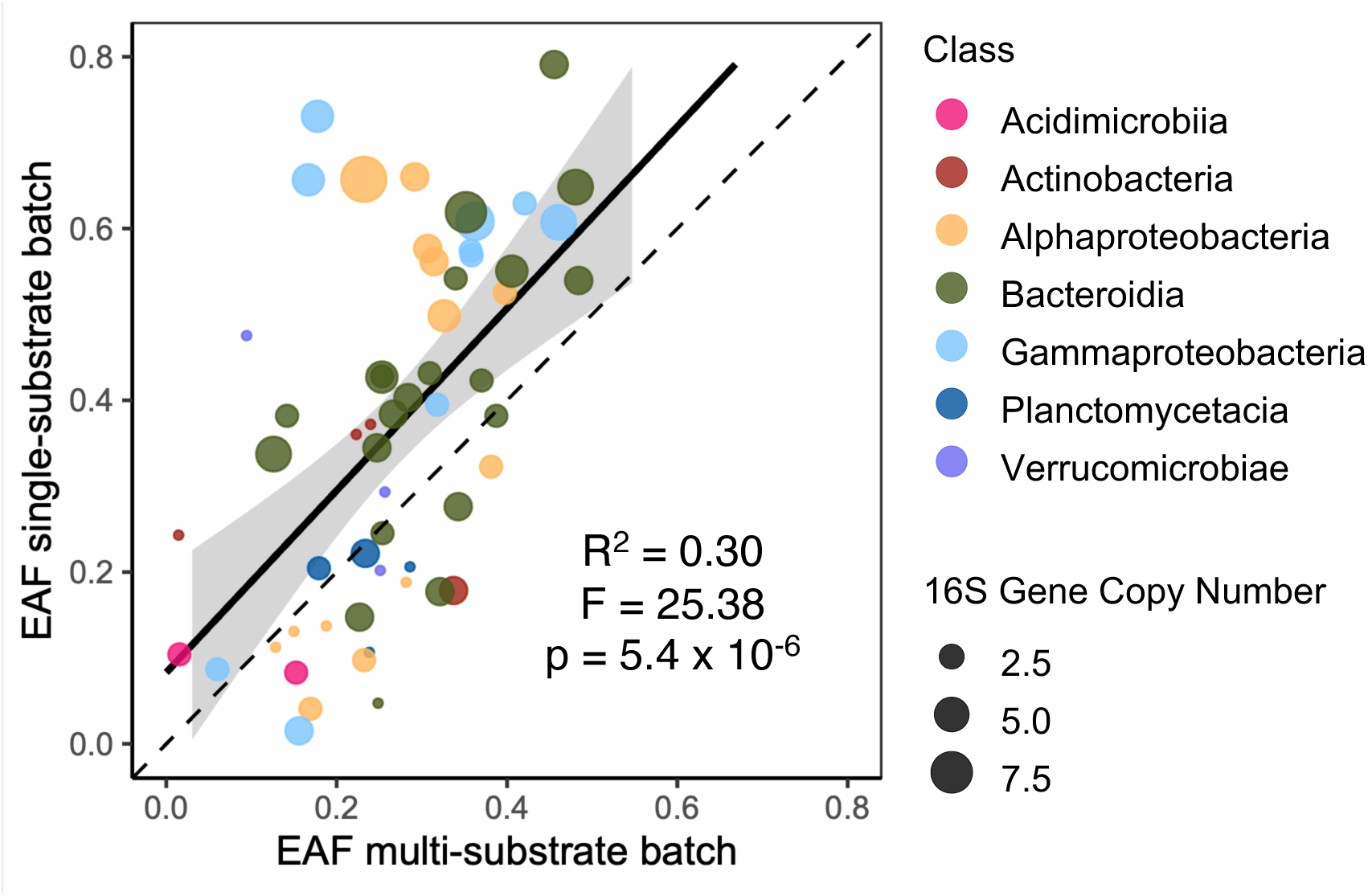
**Comparison of the excess atomic fraction (EAF) of the shared batch single-and multi-substrate incorporator ASVs**. Dashed line indicates a 1:1 line while the solid line is a linear regression with shaded area as standard error. Colors indicate the bacterial classes. Circle size represents incorporator’s 16S gene copy number. Correlation coefficients (R^2^), F-value and *p* value are from linear regression.

EAF values were significantly higher in the single-substrate than in the multi-substrate media, and higher in the multi-substrate batch than in the chemostat (Table 1). We interpret this gradient as progressive isotope dilution of the ^13^C-glucose label, and as more unlabeled substrates become available, an incorporator could draw a smaller fraction of its carbon from glucose, lowering its EAF. How much of this isotope dilution occurs could rely broadly on the bacteria’s ecological strategy – a specialist or generalist. Under the single-substrate batch experiment, copiotrophic specialists that concentrate on a single substrate as a competitive strategy have high glucose-specific EAF. With the multi-substrate chemostat, generalists that take up multiple substrates could be favored, diluting the glucose-specific EAF. This hypothesis is consistent with our multi-substrate MEP-based model, in which the short-term optimization (analogous to the multi-substrate batch) selected for substrate specialists whereas the long-term optimization (analogous to the multi-substrate chemostat) selected for substrate generalists (Vallino *et al*., 2025).

Overall, our findings demonstrate that both substrate complexity and experimental regime influence microbial metabolic activity rates and their linkage to growth strategy. These results highlight the need for caution when extrapolating qSIP-derived metabolic rates from batch systems to natural ecosystems due to potentially biased estimates of microbial heterotrophic rates towards fast growing taxa. Rates of autotrophy were not examined in this study and may be an exception due to differing metabolic constraints. A potential way to minimize bias of fast-growing taxa is to examine the unmanipulated samples at the transcriptomic level to see if the incorporators are also active *in situ*, for example (Barnett *et al*., 2021; Elkassas *et al*., 2025). Incorporating chemostat approaches and mixed-substrate designs into SIP experiments may yield more ecologically realistic estimates of microbial carbon processing and improve our ability to link microbial traits to biogeochemical functions.

## Acknowledgements

This research was supported by an NSF award DEB-1655552 (to J. A. H. and J. J. V.), and OCE-2224608 (to J.J.V.), Simons Foundation Award 549941FY22 to (J. J. V.), and the Stanley W. Watson Chair for Excellence in Oceanography (to J. A. H.). Claude (Anthropic, June 2026) was used to assist with language editing and improving the clarity of selected paragraphs. All AI-assisted text was reviewed and revised by the authors, who take full responsibility for the accuracy and integrity of the work.

---

**SI Figure 1.**
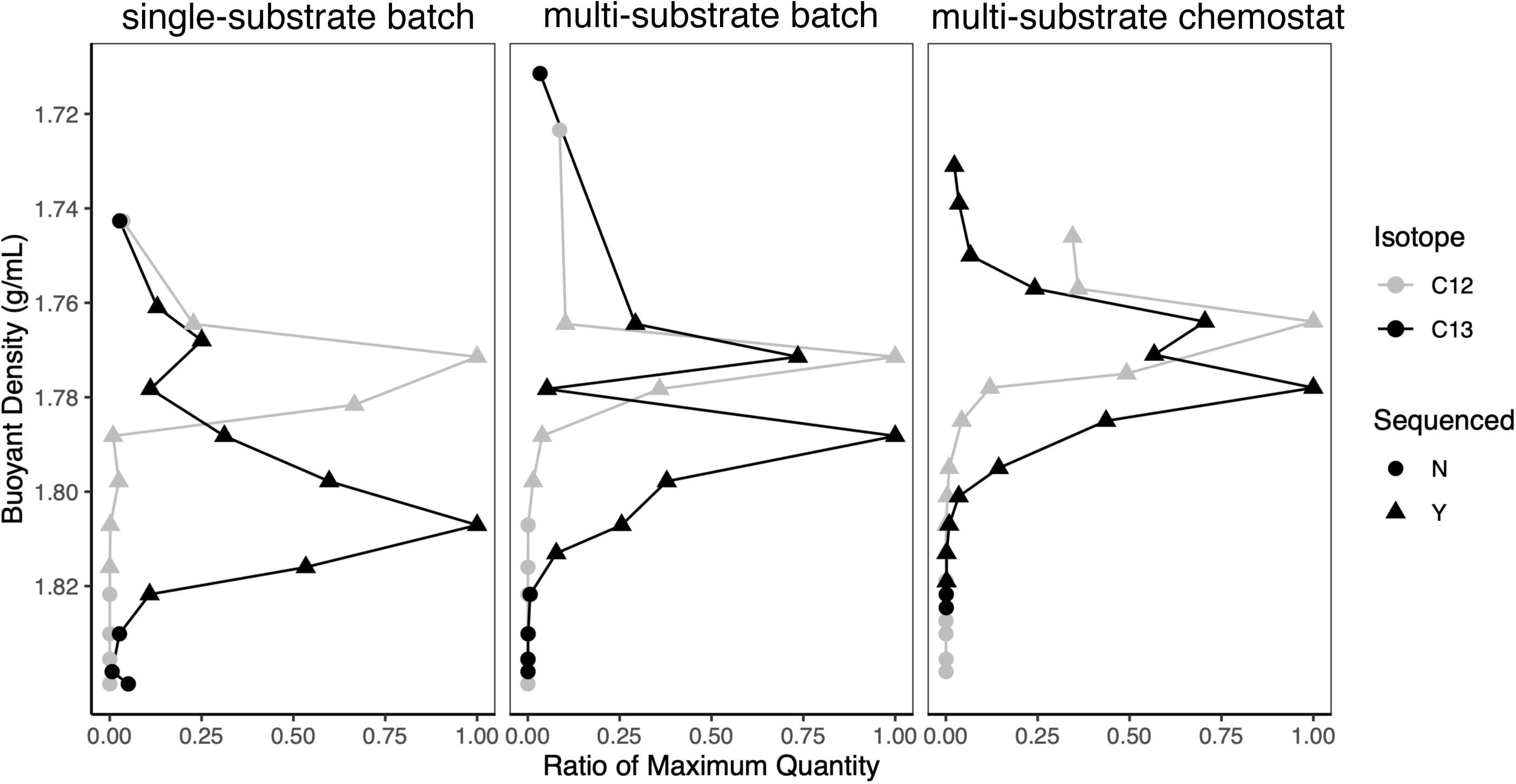
Buoyant density gradients versus ratio of maximum quantity of samples used in this study: batch single-substrate, batch multi-substrate, and chemostat multi-substrate . Colors depict the isotope enrichment (C13) or not (C12) and shape indicates whether the fraction was sent for 16S rRNA sequencing (Y) or not (N). Ratio of maximum quantity represents each fraction’s RNA concentration divided the gradient’s maximum RNA concentration.

**SI Figure 2.**
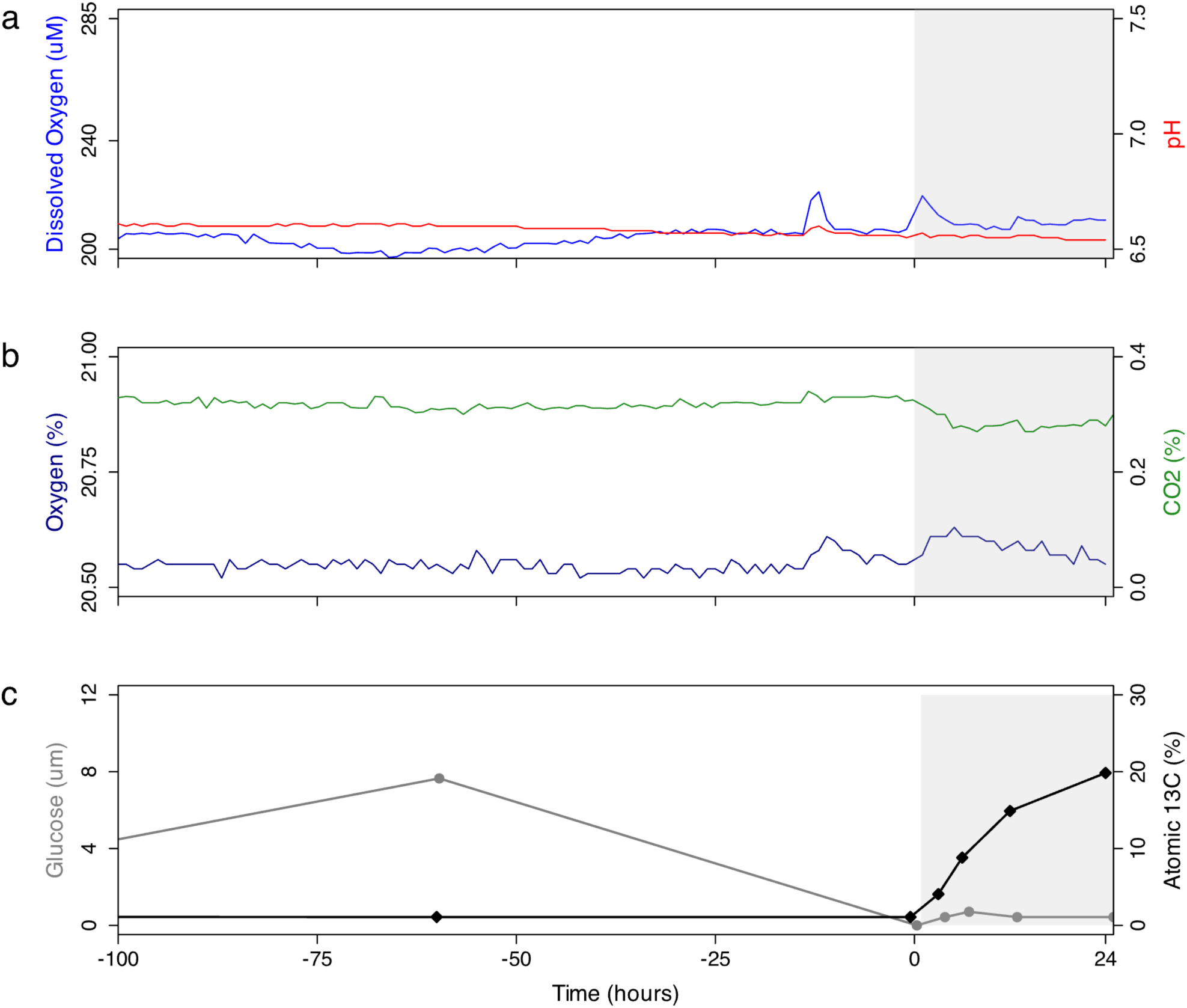
Environmental variables in chemostat during the addition of natural abundance ^12^C-control (white panel) and ^13^C-labeled treatment (gray panel). Measurements of dissolved oxygen (blue) and pH (red) (a) *in situ* and percent oxygen (blue) and carbon dioxide (green) in the chemostat headspace (b). Concentrations of (c) total glucose (gray) and atomic % ^13^C of POC (black) within the chemostat.

## References

1. Apprill, A., McNally, S., Parsons, R., and Weber, L. (2015) Minor revision to V4 region SSU rRNA 806R gene primer greatly increases detection of SAR11 bacterioplankton. Aquat Microb Ecol 75: 129–137.

2. Bailey, J.E. and Ollis, D.F. (1986) Biochemical Engineering Fundamentals, 2nd ed. New York, NY: McGraw-Hill.

3. Barnett, S.E., Youngblut, N.D., Koechli, C.N., and Buckley, D.H. (2021) Multisubstrate DNA stable isotope probing reveals guild structure of bacteria that mediate soil carbon cycling. Proc Natl Acad Sci U S A 118: e2115292118.

4. Bolyen, E., Rideout, J.R., Dillon, M.R., Bokulich, N.A., Abnet, C.C., Al-Ghalith, G.A., et al. (2019) Author Correction: Reproducible, interactive, scalable and extensible microbiome data science using QIIME 2. Nat Biotechnol 37: 1091.

5. Bulseco, A.N., Yang, W., Huber, J.A., and Vallino, J.J. (in press) Growth characteristics of natural microbial populations are skewed towards bacteria with low specific growth rates. Microbiology Spectrum.

6. Callahan, B.J., McMurdie, P.J., Rosen, M.J., Han, A.W., Johnson, A.J.A., and Holmes, S.P. (2016) DADA2: High-resolution sample inference from Illumina amplicon data. Nat Methods 13: 581–583.

7. Coskun, Ö.K., Orsi, W.D., Marshall, I.P.G., Muschler, K.A., Mitschke, N., Ferdelman, T.G., and Gomez-Saez, G.V. (2026) Hypoxia increases microbial carbon assimilation of taurine in a seasonally anoxic fjord. ISME J 20: wrag057.

8. Dal Bello, M., Lee, H., Goyal, A., and Gore, J. (2021) Resource–diversity relationships in bacterial communities reflect the network structure of microbial metabolism. Nat Ecol Evol 5: 1424–1434.

9. Dunford, E.A. and Neufeld, J.D. (2010) DNA Stable-Isotope Probing (DNA-SIP). J Vis Exp 2027.

10. Elkassas, S.M., Fortunato, C.S., Grim, S.L., Butterfield, D.A., Holden, J.F., Vallino, J.J., et al. (2025) Metabolic and population profiles of active subseafloor autotrophs in young oceanic crust at deep-sea hydrothermal vents. Appl Environ Microbiol 91: e0186825.

11. Enke, T.N., Datta, M.S., Schwartzman, J., Cermak, N., Schmitz, D., Barrere, J., et al. (2019) Modular Assembly of Polysaccharide-Degrading Marine Microbial Communities. Curr Biol 29: 1528–1535.e6.

12. Fernandez-Gonzalez, N., Huber, J.A., and Vallino, J.J. (2016) Microbial Communities Are Well Adapted to Disturbances in Energy Input. mSystems 1: e00117-16.

13. Fortunato, C.S. and Huber, J.A. (2016) Coupled RNA-SIP and metatranscriptomics of active chemolithoautotrophic communities at a deep-sea hydrothermal vent. ISME J 10: 1925–1938.

14. Gao, Y. and Wu, M. (2023) Accounting for 16S rRNA copy number prediction uncertainty and its implications in bacterial diversity analyses. ISME Commun 3: 59.

15. Gilbert, J.A., Steele, J.A., Caporaso, J.G., Steinbrück, L., Reeder, J., Temperton, B., et al. (2012) Defining seasonal marine microbial community dynamics. ISME J 6: 298–308.

16. Goldford, J.E., Lu, N., Baji, D., Sanchez-Gorostiaga, A., Segrè, D., Mehta, P., and Sanchez, A. (2018) Emergent simplicity in microbial community assembly. Science 361: 469–474.

17. Gresham, D. and Hong, J. (2015) The functional basis of adaptive evolution in chemostats. FEMS Microbiol Rev 39: 2–16.

18. Hansell, D., Carlson, C., Repeta, D., and Schlitzer, R. (2009) Dissolved organic matter in the ocean: A controversy stimulates new insights. Oceanography (Wash D C) 22: 202–211.

19. Hungate, B.A., Mau, R.L., Schwartz, E., Caporaso, J.G., Dijkstra, P., Van Gestel, N., et al. (2015) Quantitative Microbial Ecology through Stable Isotope Probing. Appl Environ Microbiol 81: 7570–7581.

20. Klappenbach, J.A., Dunbar, J.M., and Schmidt, T.M. (2000) RRNA operon copy number reflects ecological strategies of bacteria. Appl Environ Microbiol 66: 1328–1333.

21. Lueders, T. (2010) Stable Isotope Probing of Hydrocarbon-Degraders. In Handbook of Hydrocarbon and Lipid Microbiology. Berlin, Heidelberg: Springer, pp. 4011–4026.

22. Luo, E., Pham, N.D., Rogers, T.J., Sheam, M.M., Benner, B.E., Vallino, J.J., et al. (2026) Quantitative stable isotope probing (qSIP)-informed metagenomics identifies viruses infecting chemoautotrophs. Nat Commun 17: 5201.

23. Madigan, M.T., Brock, T.D., Martinko, J.M., Dunlap, P., and Clark, D.P. (2008) Brock biology of microorganisms, 12th ed. Upper Saddle River, NJ: Pearson.

24. McMurdie, P.J. and Holmes, S. (2013) phyloseq: An R Package for Reproducible Interactive Analysis and Graphics of Microbiome Census Data. PLoS One 8: e61217.

25. Mooshammer, M., Kitzinger, K., Schintlmeister, A., Ahmerkamp, S., Nielsen, J.L., Nielsen, P.H., and Wagner, M. (2021) Flow-through stable isotope probing (Flow-SIP) minimizes cross-feeding in complex microbial communities. ISME J 15: 348–353.

26. Oksanen, J. (2017) Vegan: Community Ecology Package. R package Version 2 4-3.

27. Ono, H., Tsuru, S., and Furusawa, C. (2025) Carbon source diversity shapes bacterial interspecies interactions. ISME J 19: wraf224.

28. Osburn, E.D., Weissman, J.L., Strickland, M.S., Bahram, M., Stone, B.W., and McBride, S.G. (2025) Relative abundances of bacterial phyla are strong indicators of community-scale microbial growth rates in soil. Environ Microbiome 20: 131.

29. Parada, A.E., Needham, D.M., and Fuhrman, J.A. (2016) Every base matters: assessing small subunit rRNA primers for marine microbiomes with mock communities, time series and global field samples. Environ Microbiol 18: 1403–1414.

30. Parks, D.H., Chuvochina, M., Rinke, C., Mussig, A.J., Chaumeil, P.-A., and Hugenholtz, P. (2022) GTDB: an ongoing census of bacterial and archaeal diversity through a phylogenetically consistent, rank normalized and complete genome-based taxonomy. Nucleic Acids Res 50: D785–D794.

31. Purcell, A.M., Dijkstra, P., Hungate, B.A., McMillen, K., Schwartz, E., and van Gestel, N. (2023) Rapid growth rate responses of terrestrial bacteria to field warming on the Antarctic Peninsula. ISME J 17: 2290–2302.

32. Quast, C., Pruesse, E., Yilmaz, P., Gerken, J., Schweer, T., Yarza, P., et al. (2013) The SILVA ribosomal RNA gene database project: improved data processing and web-based tools. Nucleic Acids Res 41: D590–6.

33. Reed, K., Dang, C., Walkup, J., Purcell, A., Hungate, B., and Morrissey, E. (2025) Comparing field and lab quantitative stable isotope probing for nitrogen assimilation in soil microbes. Appl Environ Microbiol 91: e0184924.

34. Roller, B.R.K. and Schmidt, T.M. (2015) The physiology and ecological implications of efficient growth. ISME J 9: 1481–1487.

35. Roller, B.R.K., Stoddard, S.F., and Schmidt, T.M. (2016) Exploiting rRNA operon copy number to investigate bacterial reproductive strategies. Nat Microbiol 1: 16160.

36. Sanz-Sáez, I., Sánchez, P., Salazar, G., Sunagawa, S., de Vargas, C., Bowler, C., et al. (2023) Top abundant deep ocean heterotrophic bacteria can be retrieved by cultivation. ISME Commun 3: 92.

37. Seemann, T. (2014) Prokka: rapid prokaryotic genome annotation. Bioinformatics 30: 2068–2069.

38. Silverman, J.D., Washburne, A.D., Mukherjee, S., and David, L.A. (2017) A phylogenetic transform enhances analysis of compositional microbiota data. Elife 6: e21887.

39. Stone, B.W., Blazewicz, S.J., Koch, B.J., Dijkstra, P., Hayer, M., Hofmockel, K.S., et al. (2023) Nutrients strengthen density dependence of per-capita growth and mortality rates in the soil bacterial community. Oecologia 201: 771–782.

40. Takahashi, S., Tomita, J., Nishioka, K., Hisada, T., and Nishijima, M. (2014) Development of a Prokaryotic Universal Primer for Simultaneous Analysis of Bacteria and Archaea Using Next-Generation Sequencing. PLoS One 9: e105592.

41. Vallino, J.J., Ahern, O., and Huber, J.A. (2025) Deriving microbial food web structure by maximizing entropy production over variable timescales. Interface Focus 15: 20250023.

42. Weissman, J.L., Hou, S., and Fuhrman, J.A. (2021) Estimating maximal microbial growth rates from cultures, metagenomes, and single cells via codon usage patterns. Proc Natl Acad Sci U S A 118: e2016810118.

43. Weissman, J.L., O’Brien, J.M., Blais, N.D., Holland-Moritz, H., Shek, K., Barbato, R.A., et al. (2026) Growth optimization predicts microbial success in a permafrost thaw experiment. Soil Biol Biochem 220: 110207.

44. Wilken, S., Yung, C.C.M., Poirier, C., Massana, R., Jimenez, V., and Worden, A.Z. (2023) Choanoflagellates alongside diverse uncultured predatory protists consume the abundant open-ocean cyanobacterium *Prochlorococcus*. Proc Natl Acad Sci U S A 120: e2302388120.

45. Youngblut, N.D., Barnett, S.E., and Buckley, D.H. (2018) HTSSIP: An R package for analysis of high throughput sequencing data from nucleic acid stable isotope probing (SIP) experiments. PLoS One 13: e0189616.

46. Zakem, E.J., McNichol, J., Weissman, J.L., Raut, Y., Xu, L., Halewood, E.R., et al. (2025) Functional biogeography of marine microbial heterotrophs. Science 388: eado5323.

47. Zheng, J., Ge, Q., Yan, Y., Zhang, X., Huang, L., and Yin, Y. (2023) dbCAN3: automated carbohydrate-active enzyme and substrate annotation. Nucleic Acids Res 51: W115–W121.

